# Imprinted antibody lineages can persist for half a century and dominate cross-neutralizing recall responses to poliovirus

**DOI:** 10.64898/2025.12.23.696092

**Authors:** Gregory C. Ippolito, Jonathan R. McDaniel, Brian C. Schanen, Diana V. Kouiavskaia, Daniel R. Boutz, Hidetaka Tanno, Megan Wirth, Erik Johnson, John J. Blazeck, Daechan Park, Bing Tan, Ellen M. Murrin, Konstantin Chumakov, Donald R. Drake, George Georgiou, Jason J. Lavinder

**Affiliations:** Texas Biomedical Research Institute and Southwest National Primate Research Center, San Antonio, TX, USA; The University of Texas at Austin, Austin, TX, USA; Sanofi, Orlando, FL, USA; FDA; Biomedicine Design, Pfizer, Cambridge, MA, USA; Houston Methodist Research Institute and Department of Pathology and Genomic Medicine, Houston Methodist Hospital, Houston, TX, USA; Cancer Immunology Project, Tokyo Metropolitan Institute of Medical Science, 2-1-6 Kamikitazawa, Setagaya-ku, Tokyo 156-8506, Japan; School of Chemical and Biomolecular Engineering, Georgia Institute of Technology, Atlanta, GA, USA; Department of Molecular Science and Technology, Advanced College of Bio-Convergence Engineering, Ajou University, Suwon 16499, South Korea; George Washington University, Washington DC, USA

## Abstract

Humoral immunity depends on a dynamic interplay between circulating polyclonal IgG antibodies and long-lived memory B cells (MBC). To elucidate this relationship, we integrated B-cell receptor sequencing with plasma IgG immunoproteomics to identify and quantify the recall response (plasma IgG and MBC) to inactivated poliovirus vaccine (IPV) in eight adults, 24–55 years after their childhood immunization. IPV immunization elicited highly skewed plasma IgG recall repertoires, dominated by a small subset (∼8%) of total antibody lineages that were detectable prior to boosting (i.e., “pre-existing”). These IgG lineages spiked logarithmically in abundance (∼10^2^-10^4^ fold-increase) and potently cross-neutralized multiple PV serotypes *in vitro* (titers ≥ 1:10^5^). Notably, these dominant plasma IgG lineages were clonally linked to class-switched MBCs that circulated in blood 21 days post-boost and bore heavily mutated VH and VL variable region genes. Importantly, IPV-primed *ex vivo* B-cell cultures (MIMIC, Modular IMmune In vitro Construct) yielded clonal expansions that mirrored the dominant post-boost plasma IgG lineages *in vivo*, providing an *in vitro* mechanistic correlate of recall immunity. Together these findings demonstrate a decades-long interconnectivity between persistent MBC clones and the serological IgG repertoire. They further highlight the central role of immunological imprinting, whereby cross-reactive MBC clones—originally primed in childhood—can be reactivated to dominate the circulating IgG recall response more than 50 years after initial exposure.

## INTRODUCTION

Historically, poliovirus (PV) has accompanied humankind through three broad phases: endemic, epidemic, and now postvaccine.^1^ The Salk inactivated poliovirus vaccine (IPV) and the Sabin attenuated oral poliovirus vaccine (OPV) instantiate one of the greatest achievements of modern vaccines in the quest for the eradication of infectious viral diseases. Since its discovery as the transmissible “filterable agent” causing paralytic poliomyelitis,^2^ and the subsequent description of its multiple viral strains,^3,4^ it has been concluded that there are three serotypes of poliovirus, designated PV I, II and III.^5^ IPV comprises three inactivated wild-type viral strains representing the serotypes Mahoney (PV I), MEF-I (PV II), and Saukett (PV III). In contrast, OPV comprises three attenuated Sabin strains of polioviruses corresponding to US reference stocks NA4 (Type 1), NB2 (Type 2), and NC2 (Type 3). Following natural exposure or vaccination, IgG becomes the predominant class of PV-specific antibody in plasma. IgG serological memory to PV may last for life and has been shown to block PV entry into the central nervous system.^4,6,7^ Historically, passive administration of gamma globulin containing anti-PV antibodies proved effective as immunotherapy against all three serotypes; however, field trials in the United States were discontinued after the introduction of Salk’s IPV in 1955 followed by Sabin’s OPV in 1961.^8,9^ These vaccines led to a rapid and exponential decline in poliomyelitis cases. By 1972, endemic transmission of wild-type poliovirus (WPV) had ceased in the United States, and by 1991, WPV was eradicated throughout the Americas.^1,10^

Early studies indicated that serological immune responses to PV infection in humans are not strictly serotype-specific.^6,11^ Additional evidence suggested that cross-protection can occur, where pre-existing antibodies against one PV serotype reduce the risk of paralytic disease (poliomyelitis) upon exposure to a different serotype.^12^ These initial observations—based on bulk serological data—were later reinforced by single B-cell cloning studies following IPV booster vaccination, which demonstrated that cross-reactive anti-PV antibodies may be more prevalent than previously recognized.^13–15^ Several important gaps remain: Is cross-reactive anti-PV IgG maintained over the long-term or only detected transiently? Does it derive from *bona fide* long-lived memory B cells (MBCs)? And critically, are these MBCs clonally linked to the long-lived plasma cells (LLPCs), that sustain antibody production and durable levels of circulating plasma antibody?

To investigate the durability of immunological memory to poliovirus, we analyzed the molecular features of the recall response to IPV boosting in healthy adults who had received their primary immunizations 24–55 years earlier during childhood in the United States. Using IgG immunoproteomics (Ig-Seq^16^) coupled with *ex vivo* activation of B cells via the Modular Immune In vitro Construct (MIMIC) system^17^ we distinguished the respective contributions of LLPCs (as proxied by secreted plasma IgG proteomics) and MBCs to the anti-PV antibody response. On average, the boosted anti-PV IgG serological repertoires were composed of approximately 70% pre-existing antibody lineages and 30% newly elicited ones, based on relative abundance by Ig-Seq. IPV boosting markedly amplified the abundance of a limited number of highly potent, mostly cross-neutralizing antibodies. These dominant plasma IgG lineages were detectable prior to boosting and were clonally related to B-cell receptors expressed by pre-existing MBCs. By coupling Ig-Seq and BCR-Seq with an antigen-primed *ex vivo* human immune model, we explicitly test—and use as *in vitro* corroboration—the predicted clonal linkage between vaccine-responsive memory B cells and the high-abundance plasma IgG lineages that dominate after boosting.

These findings demonstrate the persistence and clonal continuity of plasma IgG lineages and MBCs spanning up to ∼55 years, despite the epidemiological absence of environmental poliovirus exposure—underscoring the remarkable longevity and stability of human B-cell memory. Conceptually, these findings support a prime–boost model in which durable protection is sustained by the selective re-expansion of a small, imprinted set of cross-reactive MBC clones—an idea with practical implications for optimizing boost timing and antigen design in durable vaccines.

## RESULTS

### IPV and humoral immunological memory: multidecadal recall responses are dominated by pre-existing IgG lineages

We screened plasma from 25 adult volunteers (**Table S1**) who had received and responded to an IPV booster decades after their initial childhood vaccination (**Figure 1A**). From this group we selected eight individuals—stratified by age and neutralizing titer (**Figure 1B; Figures S1–S2**)—for detailed analysis using Ig-Seq (**Figure 1A**). Ig-Seq is a high-resolution liquid-chromatography tandem-mass spectrometry (LC-MS/MS) proteomics methodology able to identify and quantify individual antibody lineages in biological fluids following vaccination or infection.^16,18–20^ Plasma IgG was enriched by affinity chromatography against an immobilized vaccine-grade equimolar mixture of all three poliovirus serotypes (PV1, PV2, and PV3). PV-specific plasma IgG were then proteolytically digested into peptides and analyzed by LC-MS/MS. Resulting peptide spectral matches (PSMs) were mapped to a donor-specific antibody cDNA database, generated by next-generation sequencing (NGS) of peripheral blood B cells. We used heavy-chain complementarity-determining region 3 (CDR-H3) peptide sequences to quantify individual antibody lineages based on the peak intensity of their extracted ion chromatograms (XICs). Previous isobaric peptide spike-in experiments demonstrated that peak intensity correlates well with plasma antibody abundance, and that this methodology is capable of detecting antibodies circulating at low concentrations (0.4 ng/mL)^21^.

**Figure 1.**
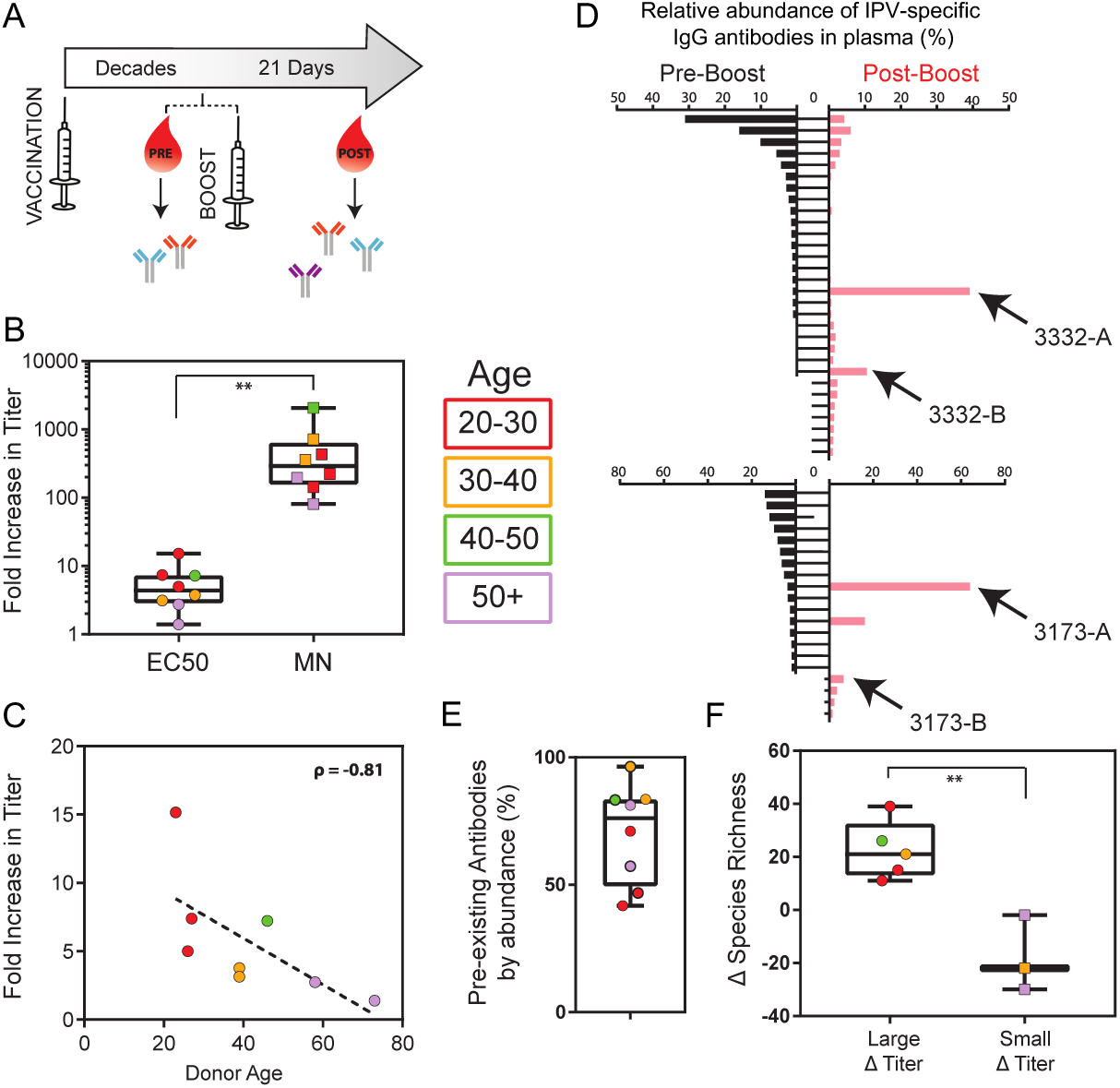
IPV elicits a recall response notable for striking increases in bulk plasma neutralization titers yet restricted expansion of only a handful of pre-existing IgG lineages. (A) Decades following pediatric vaccination, plasma and cells were isolated from adult peripheral blood donors immediately before and then 21 days following IPV boost. (B) Fold-increase in bulk plasma binding titer (EC50) against IPV antigen and fold-increase in microneutralization titer (MN) across all donors. (C) Plasma titer fold-increase (EC50) as a function of donor age (Spearman ρ = - 0.8) averaged across all three serotypes. (D) Relative abundance of IPV-specific IgG lineages pre- and post-vaccination in two representative donors 3332 and 3173. Each horizontal bar represents a single IgG lineage; labeled as pre-existing (“Pre-Boost,” black bars) or expanded/elicited (“Post-Boost,” red bars). Lineages <0.5% are not plotted for the sake of presentation. (E) Percentage of pre-existing antibodies by abundance for each donor post-vaccination. (F) Five donors with the largest fold increase in plasma titer experienced a net gain in the number of detected plasma antibody lineages, whereas the three donors with the smallest fold increase in titer experienced a net loss in detected lineages following the boost. **p < 0.05 by Mann-Whitney U test. Graphs B–F are color-coded according to donor age.

The overall magnitude of the plasma IgG response to IPV boosting, measured as the fold-increase in bulk anti-IPV IgG titer, was correlated with donor age: the two oldest donors showed the smallest increase (∼2-fold), while the three youngest donors exhibited the largest increases (∼10-fold) (**Figures 1B–1C**). Notably, however, all donors—regardless of age—experienced a striking boost in neutralizing antibody titers, ranging from 100- to 1000-fold (**Figures 1B and S2**).

To resolve the composition of these responses, we applied Ig-Seq molecular serology to identify and quantify individual anti-poliovirus IgG lineages before and after vaccination. Prior to boosting, the circulating PV-specific IgG repertoire (“pre-existing”) in each donor was diverse, comprising 21 to 118 distinct lineages. Representative examples for two donors are shown in **Figure 1D**. Post-vaccination, these pre-existing plasma IgG lineages accounted for an average of ∼70% of the total PV-specific IgG repertoire by abundance (**Figures 1E and S3**), indicating that the recall response was predominantly driven by long-lived immunological memory rather than by naïve B-cell activation. Surprisingly, only a small fraction (∼8%) of pre-existing lineages expanded significantly post-boost (defined as >3-fold relative increase and ≥0.5% absolute abundance), suggesting that a limited subset of memory-derived lineages dominate the plasma recall response.

In some cases, this dominance was extreme: a single IgG lineage accounted for 39% and 64% of the total post-vaccination PV-specific plasma IgG repertoire in donors 3332 and 3173, respectively (**Figure 1D**). Across the cohort, 7 of 8 donors exhibited at least one pre-existing IgG lineage that increased ≥50-fold in abundance post-vaccination; in 5 of these donors, that single lineage constituted ≥20% of the total post-vaccination repertoire (**Figure S3**).

IPV recall is similar to other anti-viral repertoires such as norovirus^22^, influenza^23^, and SARS-CoV-2^19^, in that the poliovirus-specific IgG repertoire post-vaccination can be highly polarized, with 4.5 ± 2.5 lineages (mean ± SD) populating the top 50% of the repertoire by abundance. Donors who exhibited the largest increases in bulk anti-IPV IgG titer also showed a corresponding rise in the number of detectable plasma antibody lineages post-vaccination. In contrast, the two oldest donors—who had the smallest increases in bulk titer—experienced a net loss in the number of detected PV-specific IgG lineages after boosting (**Figure 1F**). In contrast to seasonal influenza, which circulates perennially and elicits frequent immune reactivation, poliovirus has been largely eradicated and does not circulate in the population. Yet, despite this decades-long absence, the number of pre-existing plasma IgG lineages at the time of IPV boosting was similar between high and low responders, averaging 56 and 59 lineages, respectively.

### IPV boosts broad and potent poliovirus-neutralizing activity through the selective recall and radical expansion of a small fraction of pre-existing IgG lineages

In a subset of donors (3 out of 8), the polarization of the serological anti-PV IgG repertoire described above was extraordinarily pronounced, with just 1 to 3 lineages dominating the recall response (**Figures 1D and S3**). The intense skew toward a few dominant lineages led us to hypothesize that their selective expansion is biologically meaningful and likely plays a critical role in protective immunity.

To evaluate the functional relevance of dominant antibody responses, we recombinantly expressed the five pre-existing and two newly elicited IgG lineages that showed the greatest post-vaccination expansion in plasma abundance (∼100- to 10,000-fold increase for the pre-existing lineages). These seven monoclonal antibodies (mAbs) were tested for binding breadth and neutralization potency against all three poliovirus (PV) serotypes. Each mAb possessed a unique VH:VL pairing, with IGHV4 family gene segments utilized in 5 of the 7 mAbs. On average, the IGHV regions were heavily somatically hypermutated compared to their inferred germline segments (12.7 ± 3.1 amino acids; 14.5% ± 3.8% divergence).

Mutation frequencies in the CDR-H1 and CDR-H2 hypervariable regions were substantial (21.2% ± 8.6% and 24.3% ± 8.0%, respectively), consistent with affinity maturation through germinal center reactions. All seven mAbs bound at least one PV serotype with high affinity (EC₅₀ = 0.3–4 nM); four were cross-reactive to two serotypes, and two recognized all three (**Figure 2**). Each mAb potently neutralized at least one PV serotype *in vitro* (**Table 1**), and four cross-neutralized two serotypes with microneutralization titers ≥1:1,000.

**Figure 2.**
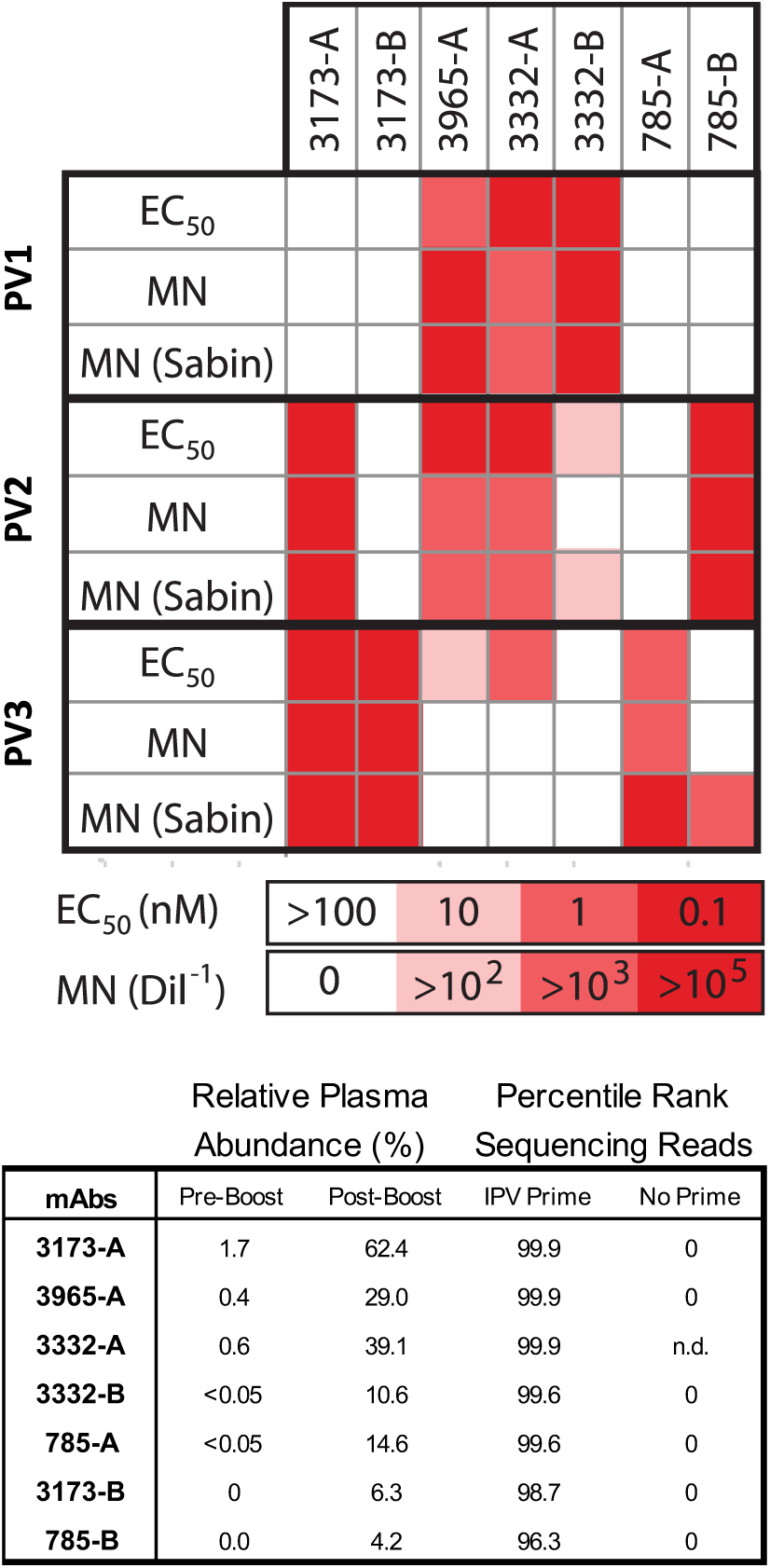
Binding (EC50) and microneutralization (MN) titers against poliovirus serotypes PV1, PV2, and PV3 for recombinant mAbs cloned from expanded B-cell clonotypes *ex vivo* which map to the most abundant pre-existing and newly elicited plasma IgG antibody lineages *in vivo* post-boost vaccination. (A) EC50 values were calculated using a four-parameter logistic model of an indirect ELISA against each serotype. Neutralization titers were determined using Salk and Sabin poliovirus microneutralization (MN) assays, as previously described ^13,24^. Numerical titers are listed in Table 1. Threshold for cross-neutralization (4 of 7 mAbs) was set at MN <10^-3^. (B) The abundance of the respective plasma IgG (pre- and post-boost) and B-cell lineages (IPV-primed and No-Prime) for each selected mAb produced recombinantly and characterized by EC50 binding and microneutralization (MN) analyses.

**Table 1.** Neutralization of Sabin and wildtype Salk polioviruses by human monoclonal antibodies mined from plasma.

| mAb ID | Serotype 1 |  | Serotype 2 |  | Serotype 3 |  |
| --- | --- | --- | --- | --- | --- | --- |
|  | Sabin 1<br>NA4 | Salk 1<br>Mahoney | Sabin 2<br>NB2 | Salk 2<br>MEF1 | Sabin 3<br>NC2 | Salk 3<br>Saukett |
| 3173-A | ND | ND | 819,200 | 409,600 | 26,214,400 | 289,600 |
| 3965-A | 19,400 | 13,700 | 9,700 | 3,400 | ND* | ND |
| 3332-A | 4525 | 1600 | 3200 | 1600 | ND | ND |
| 3332-B | 51,200 | 102,400 | 283 | ND | ND | ND |
| 785-A | ND | ND | ND | ND | 181,000 | 1,400 |
| 3173-B | ND | ND | ND | ND | 9,268,200 | 289,600 |
| 785-B | ND | ND | 4,634,100 | 4,634,100 | 1,131 | ND |

Reciprocal titer of antibody solutions at 1 mg/mL; ND, not detected.

Remarkably, the pre-existing plasma IgG lineage that dominated the post-vaccination repertoire in donor 3173 (accounting for 64% of the detected anti-PV IgG repertoire; **Figure 1D**) contained a mAb (3173-A) that cross-neutralized PV2 and PV3 with extraordinary potency. Neutralization of the Sabin PV3 strain exceeded assay limits (≥1:26,214,400; **Table 1**). MAb 3173-A also cross-neutralized both wild-type (MEF1 and Saukett) and attenuated (NB2 and NC2) PV2 and PV3 strains used in Salk IPV and Sabin OPV, respectively, with titers ranging from 1:10⁵ to 1:10⁷.

### IPV elicits long-lived memory B cells: antigen-specific, class-switched and somatically mutated MBCs can be expanded *ex vivo* and be correlated with plasma IgG molecular identity and abundance *in vivo*

A number of studies have suggested that the size of the antigen-specific plasma IgG protein repertoire in humans as well as in mice may be orders-of-magnitude smaller than the antigen-specific B-cell receptor (BCR) repertoire expressed by circulating peripheral lymphocytes.^16,21,25–27^ Additional studies have highlighted a disconnect between plasma IgG and the MBC compartment, including the observation that MBCs can persist even after plasma antibody titers have waned to near-undetectable levels.^28–31^ To better define the relationship between circulating IgG and the B-cell pool, we directly compared the abundance and neutralization activity of plasma IgG (*in vivo*) with the antigen-responsiveness of donor-matched memory B cells (*ex vivo*).

Microneutralization assays of bulk plasma collected before and after vaccination revealed a dramatic increase—ranging from 100- to 1,000-fold—in neutralization titers against all three serotypes (**Figures 1B and S2**). In contrast, bulk plasma binding titers rose only modestly (1.5- to 26-fold) over the same interval (**Figure 1B**). Due to this marked discrepancy, we hypothesized that the enhanced neutralization capacity was not due to a uniform amplification of all antibody lineages but reflected the selective expansion and skewing of a limited number of virus-neutralizing lineages. To test whether circulating MBCs) contributed to this selective serological response, we conducted parallel *ex vivo* assays measuring the response of cultured B cells—including MBCs—to PV antigens.

To measure the degree of activation and expansion in the memory compartment following antigen priming (Day 21 post-boost vaccination), we used an established *ex vivo* antigen-driven B cell activation technique, MIMIC® (Modular IMmune In vitro Construct). This immune environment—an autologous human cell-based platform—contains memory B cells, CD4+ T cells, and dendritic cells (generated from circulating monocytes) purified from PBMCs which, after 14 days of culture, recapitulates features of antigen-driven B cell stimulation following vaccination.^17,32^ MIMIC has been shown to generate antibody-secreting cells (ASCs) and IgG responses reflecting known *in vivo* immune profiles elicited by vaccines and has been used to model the humoral response to a variety of pathogens such as influenza, tetanus, hepatitis B, and yellow fever virus.^17,32–34^ Because selection, somatic hypermutation and affinity maturation do not occur in the MIMIC *ex vivo* system, ASCs that attain high abundance during the two-week culture are derived from antigen-activated MBCs that were initially seeded into the MIMIC system.

MIMIC cultures were seeded with Day 21 post-vaccination MBCs, based on reports that both “resting” and CD21^low^ “activated” MBCs at this timepoint may be recent germinal center emigrants poised to become long-lived plasma cells (LLPCs).^35–37^ Upon cognate antigen stimulation, these MBCs proliferate and differentiate into antibody-secreting cells (ASCs). To test this, each donor-derived MIMIC culture was split into a mock-treated control and an IPV-primed condition (**Figures 3A–3B**; “No Prime” and “IPV-Primed”) and maintained for 14 days. Only IPV-primed cultures generated detectable titers against PV vaccine antigens (**Figure S4**). To evaluate the degree of MBC activation, total B cells, MBCs (CD19⁺CD20⁺CD38⁻), and ASCs (CD19^low^CD20^low^CD38⁺⁺) were isolated by FACS from each culture, demonstrating that Day 21 MBCs expanded and/or differentiated in response to antigen.

**Figure 3.**
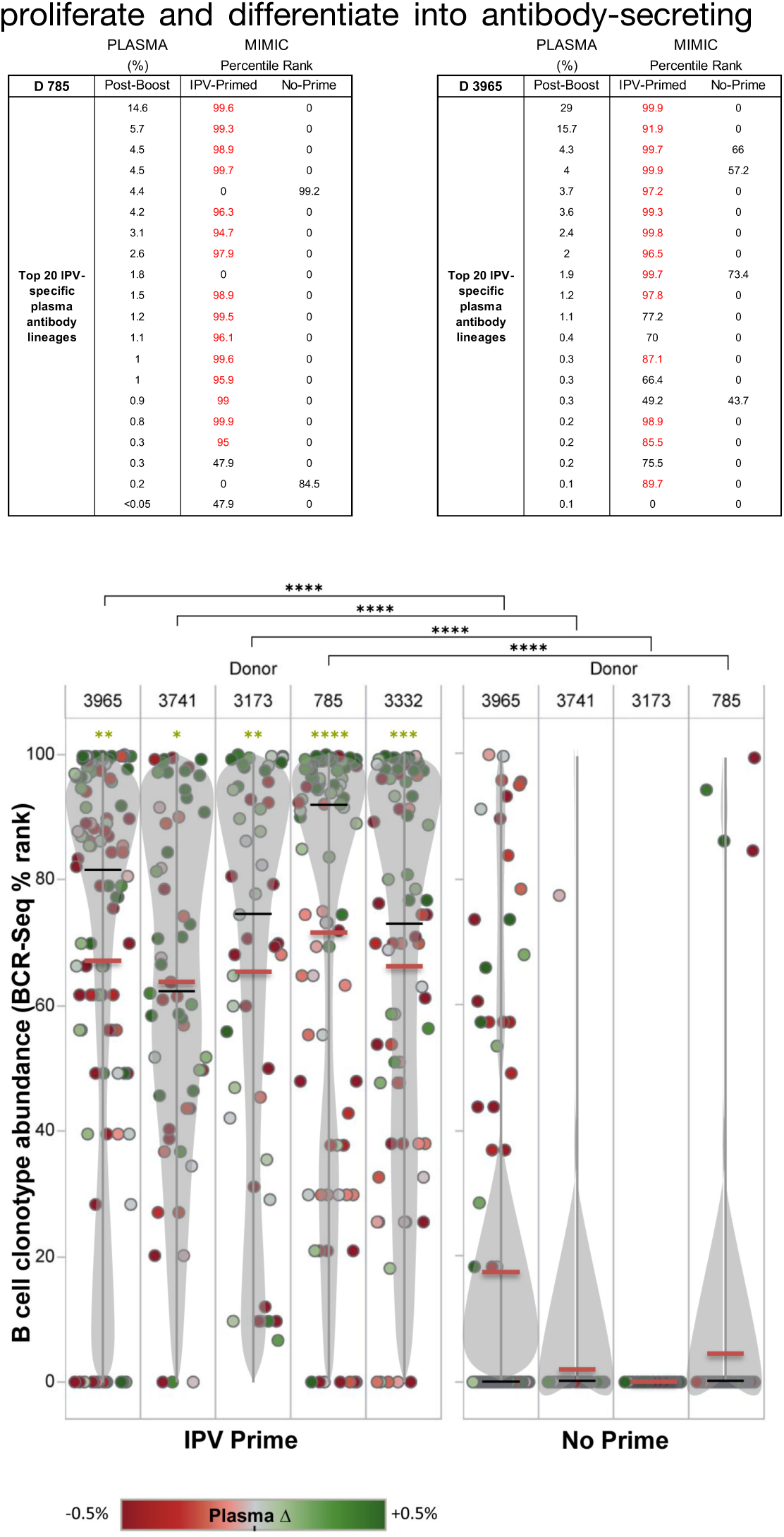
IPV elicits antigen-specific B-cell clonal expansions *ex vivo* that can be correlated with plasma IgG molecular identity and abundance *in vivo*. Predominant post-boost plasma antibody lineages derive from clonally related, peripheral blood B cells (MBCs: CD19+CD20+CD38–) expanded by antigen stimulation in the MIMIC system. **(A)**, **(B)** IgG antibody abundance pre- and post-boost (Ig-Seq) and B-cell receptor sequencing (BCR-seq) read percentile for the antigen-primed MIMIC culture (“IPV-primed”) and mock culture (“No Prime”) are shown for two donors. **(C)** For B-cell clonotypes that are detected as plasma antibody (each IgG lineage denoted as a single dot), antigen-priming of cultured donor B cells with the vaccine antigen (“IPV Primed”) caused significant expansion (higher percentile in BCR-Seq data) in all donors examined (five high titer donors shown here). Without IPV antigen priming (“No Prime”;Donor 3332 “No Prime” B-cell sequencing n.d), the majority of B-cell clonotypes detected as plasma antibody are not detected (0% rank). Bean plots (gray) represent data point density, with average and median values noted as red and black bars, respectively. Green dots represent IgG lineages with increased plasma antibody frequency from pre- to post-vaccination, whereas red dots are lineages with decreased plasma antibody frequency, with changes >0.5% or <-0.5% depicted at max color intensity. Significance (black asterisks) calculated using the Wilcoxon matched-pairs signed rank test, and linked by horizontal lines with asterisks. Gold asterisks indicate comparison of % ranks of MIMIC BCR-Seq clonotypes mapping to IgG lineages that increase (>0.5%, green) vs. lineages that decrease (<0.5%, red), calculated using the unpaired t test; ∗p < 0.05, ∗∗p < 0.01, ∗∗∗p < 0.001, ∗∗∗∗p < 0.0001.

From each donor-specific MIMIC and mock culture, total B cell receptor genes were amplified by RT-PCR, sequenced, and grouped into clonotypes. Each donor yielded approximately 2,000 to 11,000 unique BCR clonotypes. BCR-Seq of *ex vivo* MIMIC cultures revealed highly expanded clonotypes—defined by high NGS read counts—that were clonally related to plasma IgG lineages previously identified *in vivo* by Ig-Seq proteomics (**Figures 3C and S5**). These top-ranking clonotypes were consistently enriched in IPV-primed cultures compared to mock controls, particularly within the five donors exhibiting the largest fold-increases in anti-PV IgG titer (**Figures 3A–3C**). In all eight donors analyzed, the majority of PV-specific plasma antibody lineages mapped to highly ranked (>80th percentile) BCR clonotypes in IPV-primed cultures but were rarely detected in unprimed mock cultures (**Figures 3A–C and S5**).

PV-specific plasma IgG lineages that increased in abundance *in vivo* after vaccination (green circles in **Figure 3C**) were encoded by B-cell clonotypes that underwent significant expansion following antigen priming *ex vivo*, as indicated by high percentile NGS read-count ranks in “IPV-Primed”. Conversely, plasma IgG lineages that declined in abundance following booster vaccination (red circles) corresponded significantly to clonotypes showing limited IPV-primed expansion in MIMIC cultures (low percentile ranks). Notably, three donors—3965 (**Figure S3C**), 3173 (**Figure 1D**), and 785 (**Figure S3A**)—exhibited strong correspondence between anti-PV IgG abundance in plasma post-vaccination and *ex vivo* MBC expansion within IPV-primed MIMIC culture. In each of these individuals, the topmost abundant anti-PV IgG lineage, representing approximately 30%, 60%, and 15% of the post-vaccination plasma IgG repertoires, ranked 1st, 2nd, and 18th, respectively, among all B-cell clonotypes expanded in the IPV-primed MIMIC cultures (average of 6,965 clonotypes per donor). Each of these top-ranked IgG lineages was pre-existing and somatically mutated—consistent with a memory origin. Finally, we observed that B-cell expansion in response to antigen was generally lower in low-titer vaccine responders compared to high-titer responders (**Figures S5 and S6**). Together, these findings demonstrate a clear clonal link between antigen-driven B-cell expansion *ex vivo* and the increase in plasma IgG antibody abundance observed *in vivo* after vaccination.

Finally, we asked whether the clonal relationship observed between *ex vivo* MIMIC-expanded B cells and *in vivo* plasma IgG also extended to steady-state circulating MBCs *in vivo*. To test this, on a donor for which we had adequate remaining PBMC numbers, we FACS-sorted CD3⁻CD14⁻CD19⁺CD20⁺CD27⁺ MBCs from peripheral blood collected 21 days post-IPV booster (58-year-old donor D3293; **Figure S3F**). Nearly 50% of all PV-specific plasma IgG lineages in this donor had a corresponding B-cell clone within the MBC compartment. Even more striking, 12 of the 15 most abundant plasma IgG lineages were also present in the MBC pool and showed an average of ∼10% somatic mutation in their VH genes (**Table S2**). These findings firmly establish that human memory B cells are long-lived—persisting for over five decades—and are clonally linked to antibody-secreting plasma cells in the bone marrow, the experimentally established source of IgG antibodies in circulation^38,39^. This finding reinforces the classical dogma that durable humoral immunity is maintained through a clonal interconnection between memory B cells and antibody-secreting plasma cells.

## DISCUSSION

Whereas poliovirus vaccines represent one of the most successful public health interventions ever deployed against a human scourge, the mechanisms underlying their capacity to induce durable, protective humoral immunity have remained understudied and incompletely understood. Here, we employed a multi-platform approach combining IgG proteomics (Ig-Seq), B-cell receptor (BCR) deep sequencing (BCR-Seq), and *ex vivo* MIMIC modeling to elucidate how poliovirus-specific memory B cells (MBCs) and plasma IgG lineages are maintained and recalled decades after primary immunization. Our data demonstrate that inactivated poliovirus vaccine (IPV) boosters predominantly reactivate a limited number of long-lived, high-affinity, and cross-neutralizing antibody lineages originally imprinted during childhood.

Despite the extensive diversity of the circulating MBC repertoire, only a narrow subset of IgG lineages expanded in response to IPV. On average, ∼70% of the plasma IgG response post-boost was attributable to pre-existing lineages, consistent with the “Davenport phenomenon” of immunological imprinting—first described in 1956 as *persistent antibody orientation*^40^—which refers broadly to the fixating effects that initial immunization has on subsequent plasma antibody responses to antigenically distinct but related viruses. This skewed, highly polarized serological response was functionally relevant: all seven recombinantly expressed antibodies from dominant post-boost lineages neutralized at least one PV serotype, and five of seven cross-reacted and/or cross-neutralized two or more serotypes. The emergence of cross-reactive and cross-neutralizing antibodies supports the hypothesis proposed by Uhlig and Dernick^15^, in which B cells recognizing conserved viral structures are preferentially activated due to simultaneous stimulation by all three PV serotypes. This model offers a straightforward mechanism by which cross-reactive MBCs are selectively expanded and their immunoglobulins incorporated into the serological repertoire. Supporting this model, mAb 3173-A—a clonal derivative of the single most prevalent IgG lineage (64% relative abundance)—exhibited ultrapotent neutralization breadth, including what appears to be the first documented PV2:PV3 cross-neutralizing human antibody.

Building on this, a striking finding was the correlation between *in vivo* IgG antibody titers and the capacity of donor-derived peripheral-blood B cells to mount robust *ex vivo* responses in the MIMIC system. Donors whose circulating B cells yielded high-titer IgG responses in culture also displayed the highest neutralization titers in plasma post-vaccination. Moreover, the most abundant plasma IgG lineages identified by Ig-Seq were consistently clonally related to somatically mutated BCRs detected among antigen-stimulated B cells in the MIMIC cultures, often ranking in the 80+ percentile by read-count abundance. This suggests that a small number of highly functional MBC clonotypes—poised for plasma cell differentiation—dominate the recall response both *in vivo* and *ex vivo*.

Although dominant BCR clonotypes in IPV-primed cultures often mapped to known plasma IgGs, a large number of highly expanded B-cell clones could not be detected in plasma, highlighting a potential disconnect between MBC diversity and antibody secretion, as previously observed in other systems^41^. These results underscore the idea that the circulating IgG repertoire is a filtered subset of a much broader antigen-experienced B-cell pool, shaped by both transcriptional and differentiation bottlenecks.

Lastly, we provide direct evidence that the human MBC compartment retains functionally active and somatically mutated B-cell lineages for over half a century. In one 58-year-old donor, half of the circulating PV-specific plasma IgG lineages were clonally linked to MBCs isolated 21 days after IPV boosting. Among the top 15 most abundant plasma lineages, 12 were present in the MBC pool, bearing hallmark somatic mutations consistent with germinal center origin. These findings affirm the classical dogma that long-lived MBCs and long-lived plasma cells share a clonal lineage origin, and together they sustain durable humoral immunity even in the absence of antigen re-exposure.

Our study has limitations. Although our data demonstrate a tight clonal relationship between MBCs and plasma IgG lineages post-IPV boosting, we did not directly interrogate LLPCs—the bone marrow-resident producers of circulating IgG. Instead, we inferred LLPC contributions based on the detection of pre-boost plasma antibody and established fact that >90% of plasma IgG is bone-marrow LLPC-derived^30,38,39^. Furthermore, although the MIMIC system enabled robust *ex vivo* expansion of PV-specific MBCs, it cannot distinguish between two non-mutually exclusive mechanisms of recall dominance: (i) that certain MBC clones possess lower thresholds for activation and expansion, or (ii) that they are maintained at higher precursor frequencies *in vivo*. Parsing these mechanisms will require quantitative tracking of MBC precursor pools and their responsiveness *in vivo*.

An additional caveat relates to our inability to determine whether pre-existing cross-neutralizing antibodies in the two oldest donors (58 and 73 years old) arose from natural infection or pediatric vaccination. While historical context supports natural PV circulation during their early years, this confounder is absent in our youngest donors, born well after PV elimination in the U.S. Among these younger individuals, the cross-neutralizing responses we detected are most parsimoniously explained by prior IPV or OPV immunization, affirming the capacity of pediatric vaccination alone to imprint persistent and functionally diverse memory responses.

Finally, while our current study focuses on IgG, we acknowledge that IgA responses—especially mucosal—are an important component of anti-PV immunity. Substantial IgA production is known to be elicited by IPV, and future work will be needed to extend molecular serology methods to this isotype. A related priority will be the identification of viral epitopes bound by the dominant mAbs described here, particularly those that show broad neutralization. Epitope mapping—particularly within the context of the poliovirus capsid and the sterically shielded “canyon” region surrounding the virion’s icosahedral fivefold axis of symmetry^42,43^—may help delineate structural determinants of broad protection. Notably, the CDR-H3 loops of our most potent antibodies (e.g., mAb 3173-A) were relatively short (11–14 amino acids), in contrast to previously described cross-neutralizing mAbs derived from B cells, which exhibited longer CDR-H3 regions (17–26 residues)^14,44^. This observation raises the hypothesis that shorter CDR-H3s may better accommodate access to epitopes flanking the canyon, whereas longer loops may be necessary to penetrate more deeply into the canyon and interfere with poliovirus receptor CD155 engagement.

This study reveals a mechanistic basis for how IPV induces lifelong protective immunity to poliovirus: through the selective expansion of a small cadre of potent, cross-reactive MBCs imprinted during early childhood, which manifests serologically as a skewed dominant abundance of as few as one to three IgG lineages in IPV post-boost plasma. These findings suggests that vaccine protection is derived from the dynamic interplay between cross-serotype neutralizing MBC and circulating IgG. We propose that the high abundance of any given IgG lineage serves as *prima facie* evidence of its immunological relevance—a principle broadly applicable across vaccines—as exemplified by donor D3173, whose dominant lineage accounted for 64% of the post-boost IgG repertoire and potently cross-neutralized PV2 and PV3. This result supports the prioritization of strategies that intentionally prime and recall broadly neutralizing memory clones, rather than indiscriminate repertoire broadening, to maximize durability and cross-serotype protection. As global efforts to eliminate poliovirus enter their final phase, these insights may inform the design of next-generation vaccines targeting other persistent viral pathogens—particularly those that rely on the durable recall of broadly neutralizing antibody responses.

## METHODS

### Human donors and PBMC isolation

PBMCs used in the assays were acquired from normal healthy donors who provided informed consent and were enrolled in a Sanofi apheresis study program (protocol CRRI 0906009). Blood collections were performed at OneBlood (Orlando, FL) using standard techniques approved by their institutional review board. Within hours following their harvest from the donor, the enriched leukocytes were centrifuged over a ficoll-paque PLUS (GE Healthcare, Piscataway, NJ) density gradient as described previously ^45,46^. PBMCs at the interface were collected, washed, and cryopreserved in IMDM media (Lonza, Walkersville, MD) containing autologous plasma and DMSO (Sigma–Aldrich, St. Louis, MO). Subject cohort is detailed in Table S1.

### Antigen priming of donor B cells

Donor B cell responses to the inactivated polio vaccine (IPV) were generated using the MIMIC B-cell LTE module as described ^33^. The modules were prepared from aphaeresis samples collected 21 days post-vaccination with IPV. The module consisted of autologous magnetically purified B-cells, CD4+ T cells, and dendritic cells (DCs) that had been primed with IPV, or in the case of the mock controls lacked IPV-priming. Briefly, the DCs were generated by culturing monocytes positively selected using magnetic beads to CD14 (Stem Cell). The cells were then cultured for 7 days in X-VIVO 15 (Lonza) plasma-free media supplemented with GM-CSF (R&D Systems, Minneapolis, MN) and IL-4 (R&D Systems). Following IPV priming, the DCs were co-cultured with magnetically isolated CD4 T cells (Stem Cell). Total untouched B cells were then magnetically isolated by negative selection (Stem Cell EasySep^TM^ Human B Cell Isolation Kit) and primed with IPV (mock B cells were not antigen-primed). Following IPV priming, the B cells were harvested, washed and co-cultured with the DC-T cell cultures for 14 days.

The MIMIC B-cell modules were harvested on day 14 and centrifuged at 3000xg for 7 minutes at 4°C. The cell pellets were washed several times prior to cell sorting by FACS (Bio-Rad S3^TM^). Cells were sorted into memory (CD19+CD20+CD38–) and antibody-secreting (CD19^lo^CD20^lo^CD38++) cell populations and prepared in RLT RNA lysis and stabilization buffer (Qiagen) encompassing unsorted, memory, and plasmablast cells. Donor 3173 MIMIC B-cells were also provided as viably frozen cells for high throughput VH:VL pairing as previously described ^47^.

### BCR-Seq

Next-generation sequencing of donor antigen-primed and mock B-cell modules using Illumina 2×300 MiSeq was performed and processed as previously described ^21,48^.

### Ig-Seq

Ig-Seq analysis has been previously described.^21,49^ Briefly, in this study, for each pre-vaccination and post-vaccination plasma sample, total IgG was purified from 4 mL of plasma using the Melon Gel IgG purification kit (Pierce) according to the manufacturer’s protocol with the following modifications: for each 4 mL of plasma, a gravity column was constructed containing 2 mL of settled Melon Gel resin. Plasma was diluted 1:10 in Melon Gel Purification Buffer, applied to the column, and the IgG-containing flowthrough was collected and reapplied to the regenerated column a second time to ensure high IgG purity of the Melon Gel column flowthrough.

The purified plasma IgG was dialyzed into 20 mM sodium acetate, pH 4.5 and concentrated in a Vivaspin Turbo 15 (30K MWCO) (Sartorius) to 10 mg/mL. The IgG was then digested with 1 mL immobilized pepsin resin (Pierce) per 10 mg of IgG. Pepsin digestion was allowed to proceed for three hours, shaking vigorously at 37 °C. After the pepsin digestion was complete, the digestion slurry was spun through an Ultrafree-MC centrifugal filter unit (EMD Millipore) at 4000xg and the F(ab’)2-containing filtrate aspirated into a separate eppendorf. The resin (still in filter unit) was vortexed with 1 volume of 2X PBS and wash filtrate collected by spinning at 4000xg, with three resin washes total. Original filtrate and three wash filtrate were combined and pH adjusted to 7 with Tris buffer.

Vaccine-grade poliovirus serotypes 1, 2, and 3 (PV1, PV2, and PV3) were generously provided by Sanofi Pasteur (Marcy L’Etoile, France) at vaccine-ready formulation and concentration. To increase solubility of PV1 and PV3 during concentration of the inactivated virus, the NaCl concentration was adjusted to a final concentration of 0.6 M and buffered with 100 mM Tris, pH 8.5. Each serotype was then concentrated in a VivaSpin Turbo 15 (10K MWCO) (Sartorius) to a final concentration of 2 mg/mL as determined by absorbance at 268 nm (1 mg/mL poliovirus = OD268 of 7.7). Concentrated poliovirus serotypes were then each injected into 30K MWCO Slide-a-Lyzer cassettes (Pierce) and dialyzed overnight at 4°C into PBS with NaCl concentration increased to 0.6 M. Affinity chromatography for the isolation of antigen-specific F(ab’)2 was carried out by first individually coupling 1 mg of each vaccine-grade poliovirus serotype onto 0.2 g of dry N-hydroxysuccinimide (NHS)-activated agarose (Pierce) by overnight rotation at 4 °C. The coupled agarose beads were washed with PBS and unreacted NHS groups were blocked with 1 M ethanolamine, pH 8 for 20 min at room temperature, washed with PBS, and packed into a 5 mL chromatography column (Pierce). The column was then washed with 5 volumes of 1 M NaCl to elute non-specifically bound (unconjugated) antigen and then 10 volumes of PBS to equilibrate. F(ab’)2 fragments were applied to the antigen affinity column in gravity mode. The column was subsequently washed with 20 volumes of PBS, 10 volumes of ddH_2_0, and eluted into Maxymum Recovery 1.5 mL tubes (Axygen Scientific, CA, USA) using 1 mL fractions of 50 mM HCl. The flowthrough, wash, and each 1 mL elution fraction (immediately neutralized with NaOH/Tris) were analyzed by indirect ELISA against vaccine-grade poliovirus. Elution fractions showing an ELISA signal were combined and concentrated under vacuum to <0.5 mL volume and the combined, concentrated affinity column elution was desalted into 10 mM Tris, pH 8.0 using two or three 0.5 mL 45K MWCO Zeba spin columns (Pierce). The desalted elution was further concentrated under vacuum to ∼0.5 mg/mL in a Maxymum Recovery 1.5 mL tube. Ig-Seq proteomic sample preparation and LC-MS/MS analysis was then performed as previously described ^21^.

### Microneutralization assay

Poliovirus microneutralization assays were performed on the bulk plasma and recombinant mAbs according to the WHO protocol as described ^13,24,50^.

### Antibody screening and expression

Antibodies were selected based upon lineage rank in the post-vaccination plasma Ig-Seq analyses for each of the four donors with the highest neutralization titers against the IPV antigen. The exact lineage member chosen for gene synthesis (IDT) was selected based upon the presence and quantity of corresponding peptides in the Ig-Seq analysis, with the highest ranked CDR-H3 peptide and then full length (using upstream VH peptide sequences) amino acid sequence present within the Ig-Seq data selected. For the VL, either VH:VL paired amplicon 2×300 MiSeq was used as described here ^47^ to select the corresponding light chain sequence (for donor 3173 only) or it was identified via phage scFv screening (e.g. no panning required) on each synthesized VH produced as a VL shuffle library or, if necessary, 1–2 rounds of phage panning as described ^51^.

For expression of full length recombinant mAbs, the identified VH and VL were each cloned into pcDNA3.4 as described^52^, expressed in HEK293F cells, and purified over protein G gravity flow columns. Purified monoclonal antibody was then analyzed by indirect ELISA against vaccine grade PV1, PV2, and PV3 and EC50 values fit using four-parameter logistic (4PL) curves in GraphPad Prism software.

## DATA AVAILABILITY

Sequence data are openly available in a public repository. Raw Illumina sequencing reads for B cell receptor sequences (MIMIC BCR-Seq) have been deposited at NCBI Sequence Read Archive (SRA) under BioProject ID: PRJNA1392832 and are publicly available as of the date of publication. All other data are presented in the figures and supplementary materials.

## AUTHOR CONTRIBUTIONS

Conceptualization: G.C.I., D.R.D. III, G.G., and J.J.L.; Methodology: G.C.I., J.R.M., B.C.S., D.R.B. and J.J.L.; Investigation: J.R.M., B.C.S. D.V.K., D.R.B., H.T., E.J., M.W., J.J.B., B.T., E.M.M., and J.J.L.; Formal Analysis: G.C.I., J.R.M., B.C.S. D.V.K., D.R.B., D.P, and J.J.L.; Writing (original draft): G.C.I., J.R.M., and J.J.L.; Writing (review and editing): G.C.I., J.R.M., E.J., K.C., G.G., and J.J.L.

## ACKNOWLEDGEMENTS

Next-generation sequencing was performed by the Genomic Sequencing and Analysis Facility at UT Austin, Center for Biomedical Research Support, RRID#: SCR_021713. This work was supported by the Defense Threat Reduction Agency (HDTRA1-12-C-0105).

## Supplemental Figures

**Figure S1:**
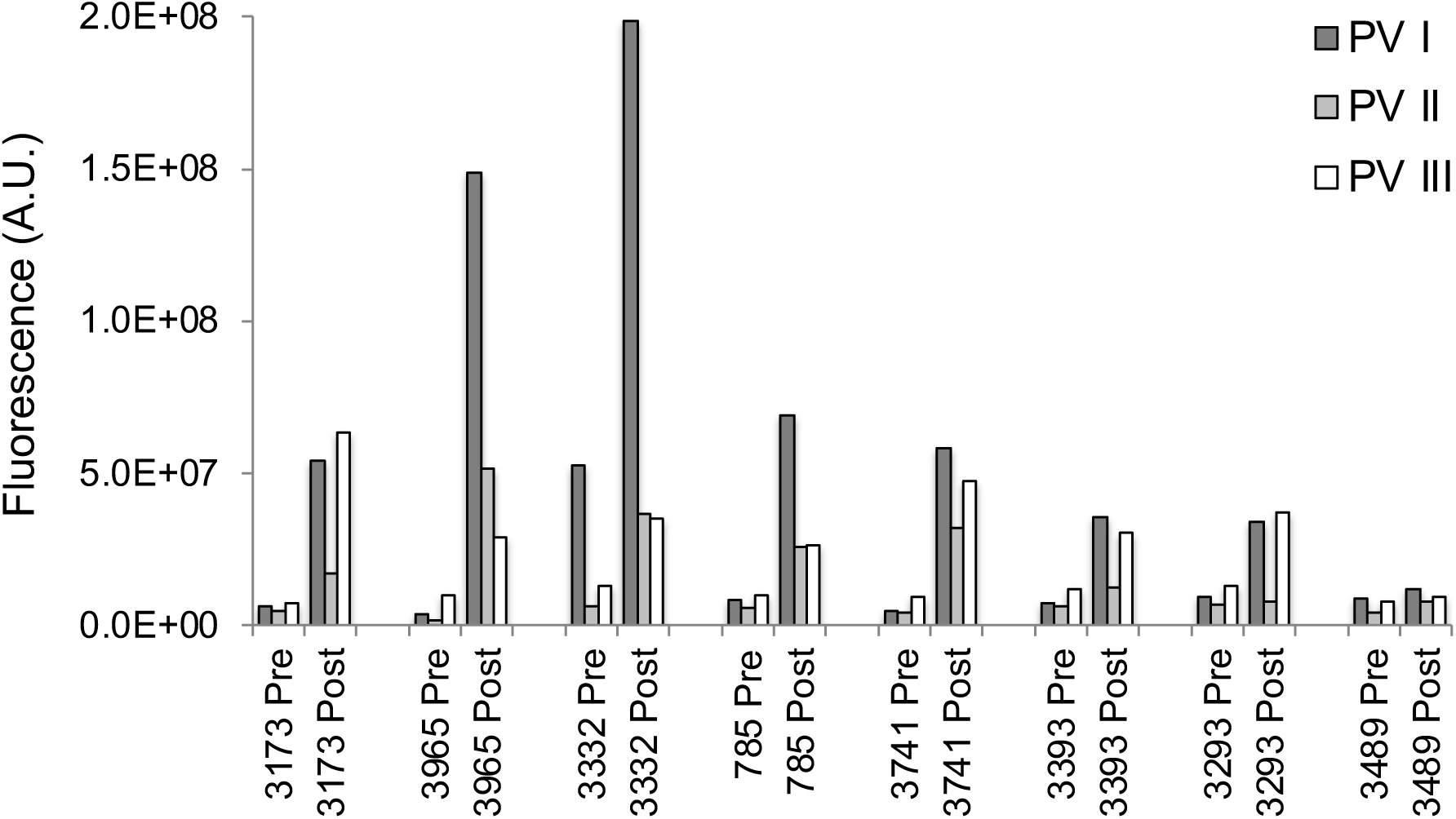
Eight IPV-immunized donors were prioritized for study according to fluorescence adherence inhibition assay of vaccine-induced plasma antibodies. Fluorescence adherence inhibition assay (fADI), using inactivated polio virus labeled with fluorescent secondary antibodies, was used in a high-throughput Luminex bead-based assay to prioritize vaccinated subjects for study according to the quantitative ability of plasma antibodies to prevent adherence of labeled virus to target Vero cells ^53^. PV1 = Mahoney; PV2 = MEF1; PV3 = Saukett.

**Figure S2:**
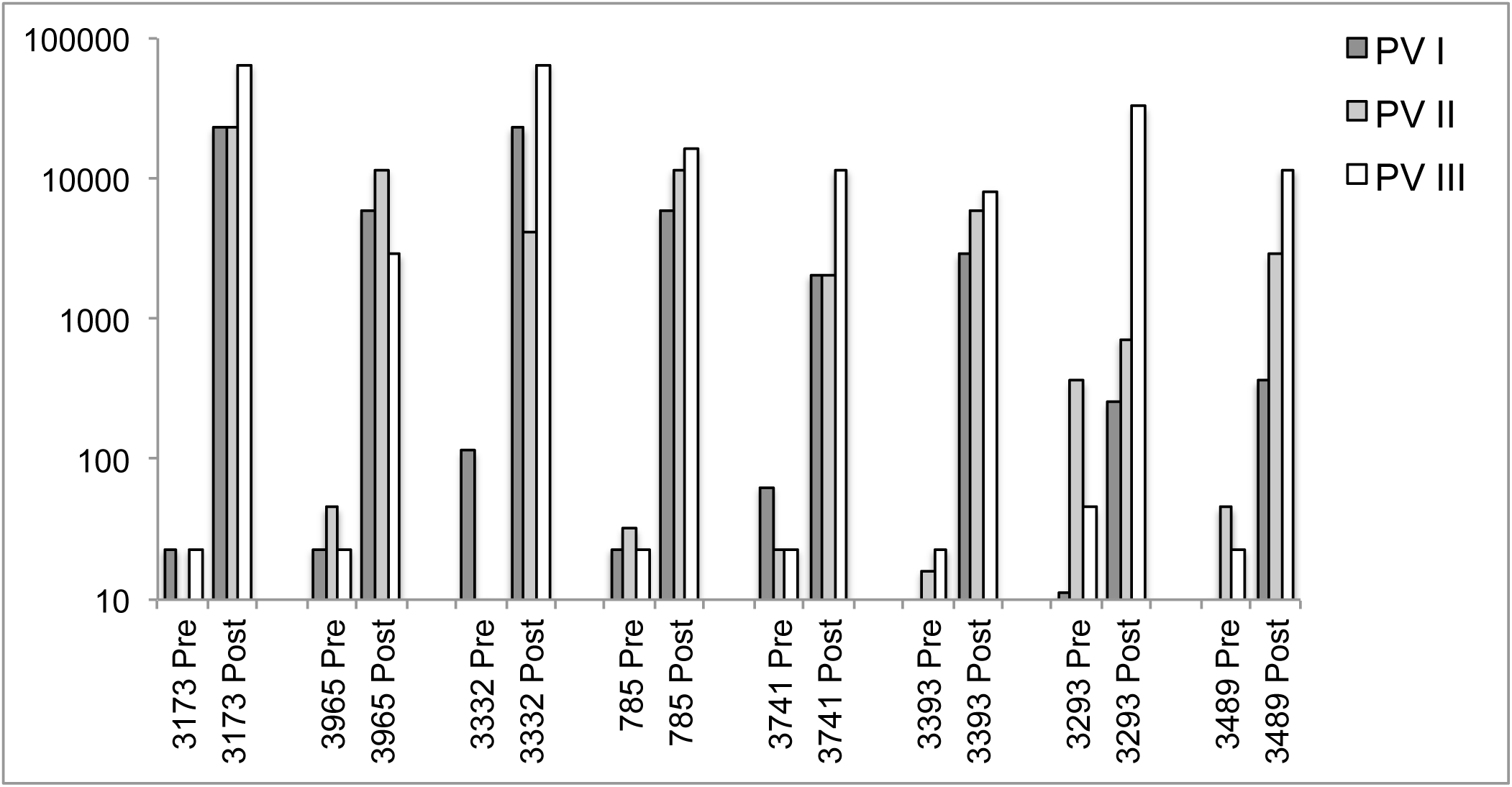
Plasma neutralization titers for the selected IPV-immunized study subjects. The pre-vaccination neutralization titers for all eight study subjects are above the baseline cutoff for protection (1:8) for each poliovirus serotype. Neutralization titers are listed here as plasma dilution, as determined by the poliovirus microneutralization assay using IPV antigen. PV1 = Mahoney; PV2 = MEF1; PV3 = Saukett.

**Figure S3:**
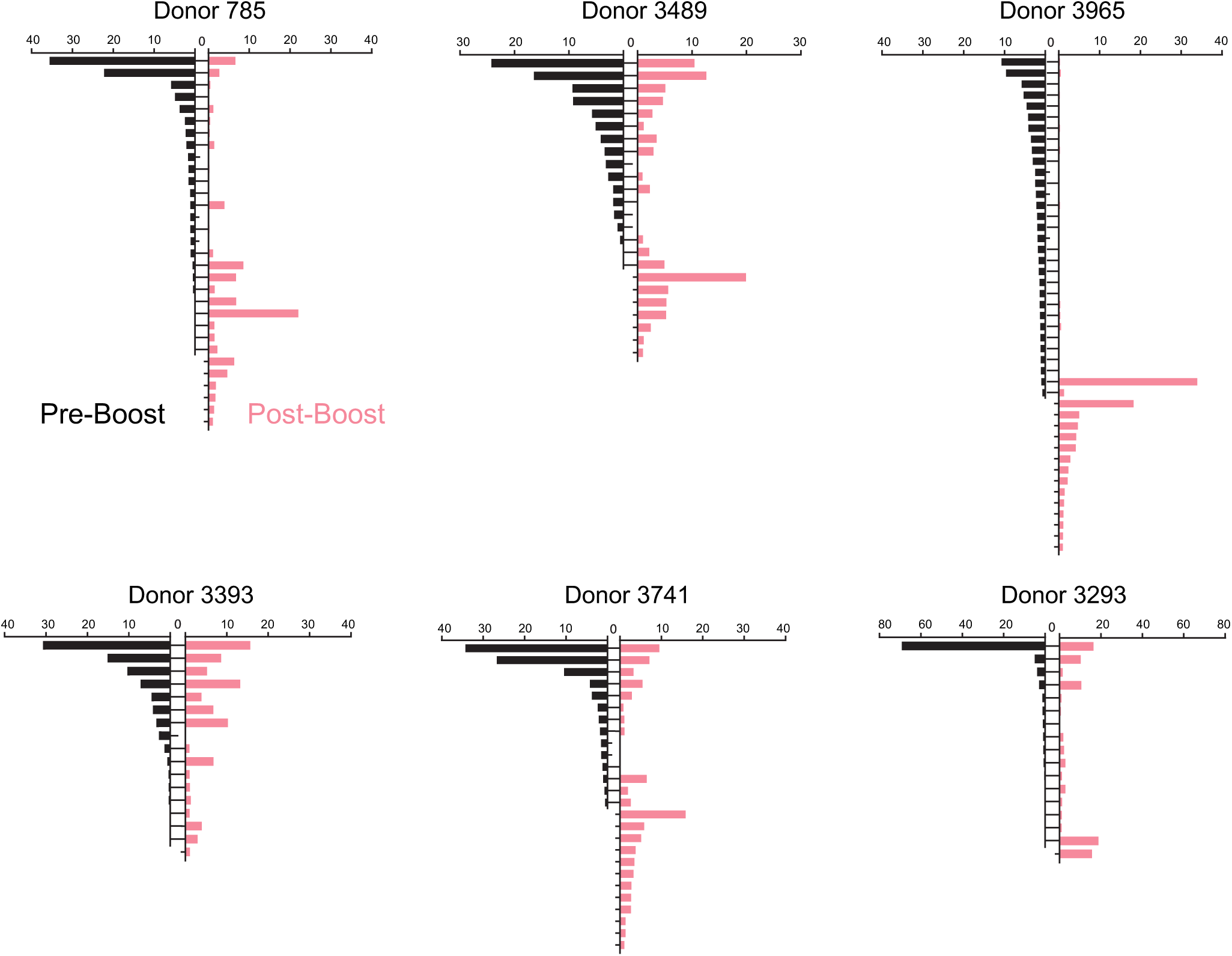
Relative percentage abundance of IPV-specific IgG lineages pre- and post-vaccination. Each horizontal bar represents a single antibody lineage. Antibodies are labeled as pre-existing (“Pre-Boost,” black bars) or newly elicited (“Post-Boost,” red bars). Lineages <0.5% are not plotted for the sake of presentation.

**Figure S4:**
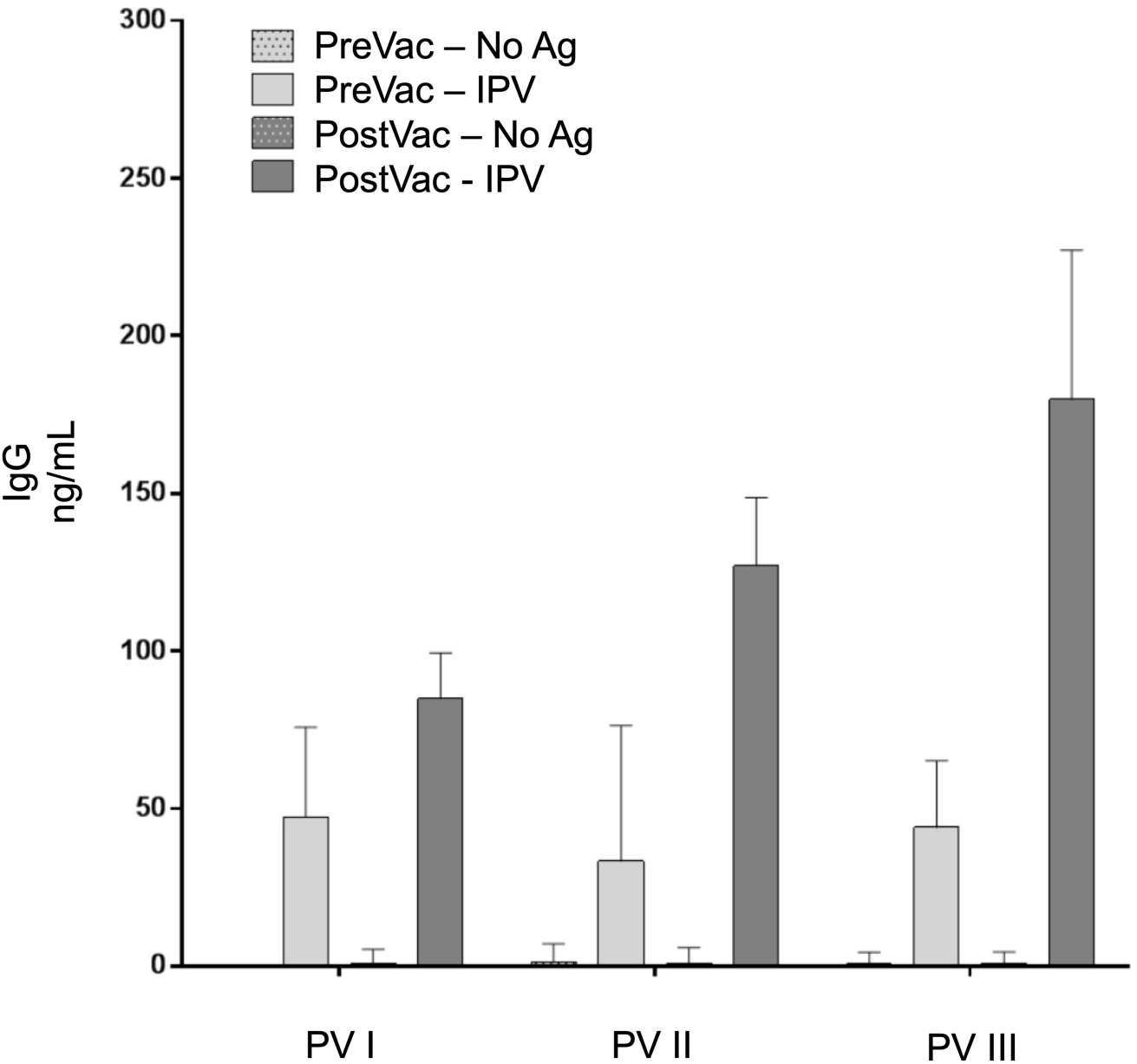
Average donor polio-specific IgG responses in MIMIC cultures. Cultures were established using donor PBMC samples (pre-/post-vaccination). Each analysis included no-treatment (No Ag) and IPV treatments. The average responses ± SD against each of the three poliovirus serotypes is shown. PV1 = Mahoney; PV2 = MEF1; PV3 = Saukett.

**Figure S5:**
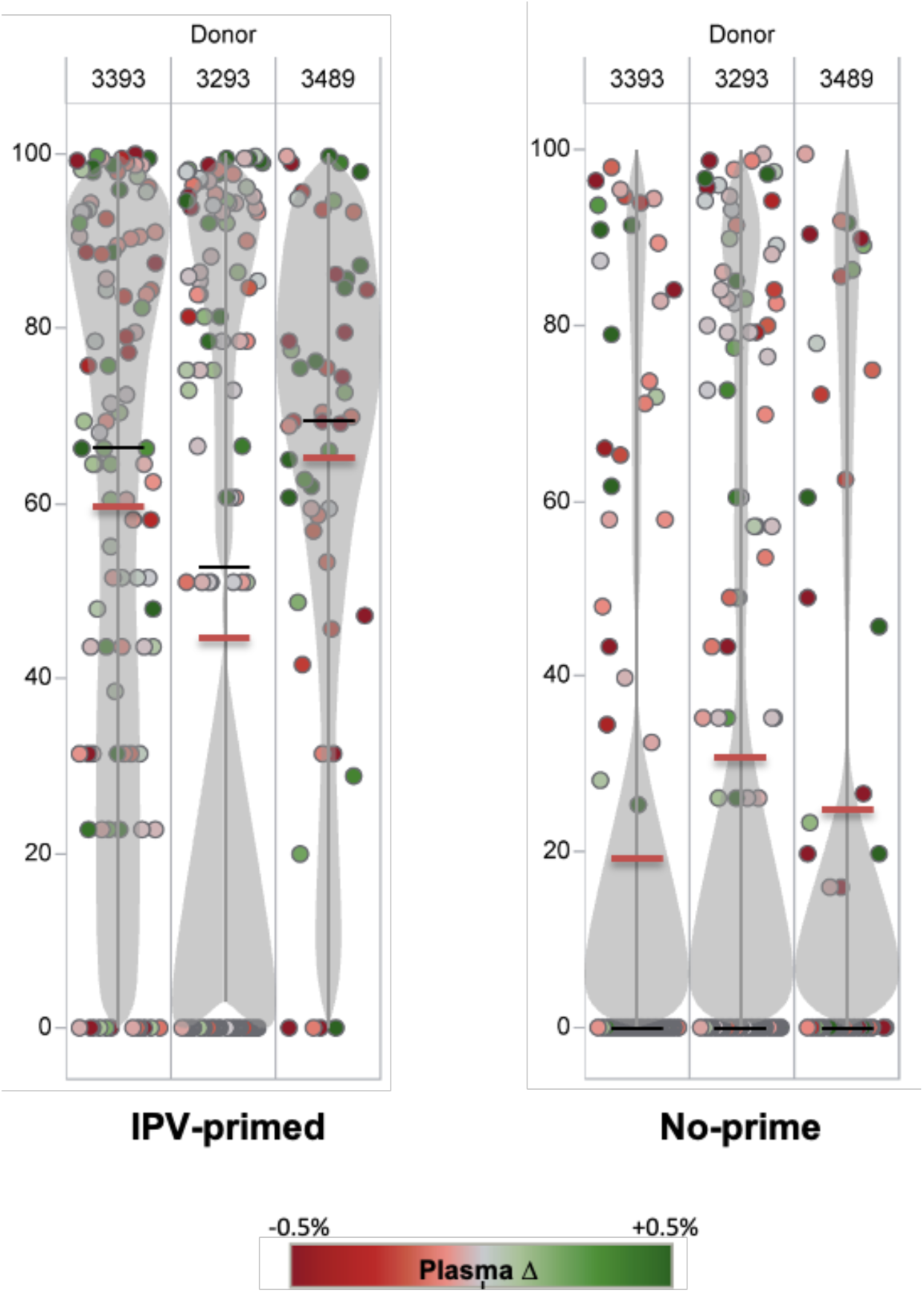
BCR-Seq profile of the MIMIC lymphocyte modules (antigen-primed and negative no prime control) for antibody lineages detected in the plasma of three low titer responders. For B cell lineages that are detected as plasma antibody (each lineage denoted as a single dot), antigen-priming of cultured donor B cells with the vaccine antigen IPV causes significant expansion (higher % rank in BCR-Seq data) in all donors examined. Bean plots (gray) represent population density, with average and median values noted as red and black bars respectively. Green-filled dots represent lineages with increased plasma antibody frequency after vaccination, whereas red dots are lineages with decreased plasma antibody frequency, with changes above or below 0.5% at max color intensity. Without the IPV antigen prime (e.g. no prime control), the B cell lineages detected as plasma antibody are mostly very low frequency within the negative control lymphocyte module, many of them undetectable by BCR-Seq.

**Figure S6:**
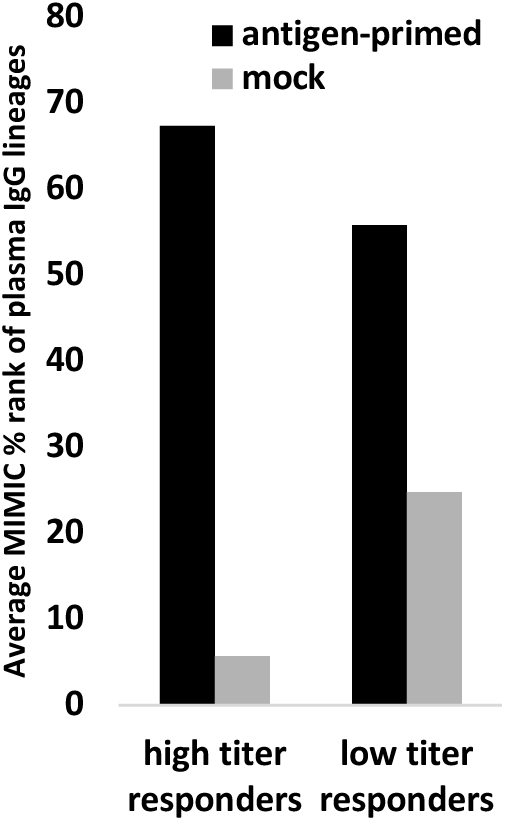
Ex vivo (MIMIC vs mock) expansion of B cell clonotypes detected as PV-specific plasma IgG lineages in high titer vs. low titer donors. Antigen-priming of cultured donor B cells with the vaccine antigen IPV (i.e. MIMIC cultures) induces significant expansion (higher % rank in MIMIC BCR-Seq data) of B cell clonotypes that are detected as plasma IgG as compared to a mock culture without antigen priming. This is seen in all seven donors examined (donor 3332 mock data not available). However, the effect is significantly increased in the higher titer responders (top four) as compared to the three low titer responders (3393, 3293, and 3489).

## Supplemental Tables

**Table S1:** Donor profile.

| ID | Age | Sex | BloodType | HLA | CMV | Vaccine | Date |
| --- | --- | --- | --- | --- | --- | --- | --- |
| 259 | 29 | M | O+ | A*0101 A*2902 B*4403 B*5701 | positive | Polio | 12/3/12 |
| 720 | 50 | M | A+ | A*0101/01n A*6801 B*0801 B*1518 | positive | Polio | 12/17/12 |
| 785 | 26 | M | B+ | A*1 A*- B*37 B*44 | negative | Polio | 12/3/12 |
| 1529 | 28 | M | A+ |  | TBD | Polio | 1/12/12 |
| 2309 | 52 | M | A+ |  |  | Polio | 12/3/12 |
| 2454 | 23 | F | AB+ |  | negative | Flu (Injections) | 9/12/13 |
| 2464 | 28 | M | O+ |  | negative | Polio | 12/13/12 |
| 2551 | 33 | M | O+ |  | TBD | Flu (Injections) | 9/23/13 |
| 3085 | 45 | F | O+ |  | positive | Polio | 1/4/12 |
| 3173 | 46 | M | A- |  | negative | Flu (Injections) | 10/9/13 |
| 3292 | 52 | M | O+ |  | TBD | Polio | 6/11/12 |
| 3293 | 58 | M | A+ |  | TBD | Polio | 6/4/12 |
| 3332 | 39 | M | O- |  | TBD | Flu (Injections) | 10/28/13 |
| 3393 | 39 | F | A+ |  | TBD | Flu (Injections) | 9/19/13 |
| 3458 | 25 | F | O- |  | TBD | Flu (Injections) | 9/16/13 |
| 3489 | 73 | M | A+ |  | positive | Polio | 11/27/12 |
| 3580 | 61 | F | O+ |  | positive | Polio | 7/16/12 |
| 3706 | 48 | M | A+ |  | positive | Polio | 12/4/12 |
| 3708 | 47 | F | O+ |  | positive | Flu (Injections) | 9/19/13 |
| 3741 | 27 | F | A+ |  | positive | Flu (Injections) | 10/4/13 |
| 3780 | 65 | M | B- |  | negative | Polio | 12/20/12 |
| 3965 | 23 | M | O+ |  | negative | Polio | 12/18/12 |
| 4097 | 40 | M | A+ |  | positive | Flu (Injections) | 9/30/13 |
| 4177 | 38 | F | A+ |  | negative | Flu (Injections) | 11/5/13 |
| 4260 | 49 | M | A+ |  | positive | Polio | 1984, 2000, 12/10/2012 |

**Table S2:** Top 15 plasma IgG lineages. Plasma IgG abundance and peripheral memory B cell (MBC) detection by NGS at 21 days post-vaccination. Donor D3293 (58 yr).

| Antibody lineage<br>ID | Plasma IgG |  | Peripheral MBC |  |
| --- | --- | --- | --- | --- |
|  | Pre-vaccination<br>(%) | Post-vaccination<br>(%) | Day 21 mBC<br>(present in NGS?) | SHM<br>(% nt) |
| 7520 | 0.003 | 17.0 | + | 11.5 |
| 12154 | 57.2 | 14.9 | + | 10.6 |
| 10081 | 0 | 14.3 | + | 8.8 |
| 693 | 2.5 | 9.6 |  | 4 |
| 16733 | 4.1 | 9.3 |  | 3.4 |
| 1328 | 0.6 | 2.6 | + | 5.5 |
| 3855 | 0.3 | 2.6 | + | 3.1 |
| 8189 | 0.8 | 2.0 | + | 12.2 |
| 2027 | 0.8 | 1.7 | + | 15.9 |
| 3871 | 3.2 | 1.4 | + | 7 |
| 2750 | 0.3 | 1.1 | + | 5.2 |
| 10468 | 0.4 | 1.0 | + | 2.4 |
| 13407 | 0.2 | 1.0 | + | 7 |
| 10403 | 0.1 | 0.9 |  | 6.3 |
| 646 | 1.1 | 0.9 | + | 5.1 |

